# Topological demarcation by HMGB2 is disrupted early upon senescence entry across cell types and induces CTCF clustering

**DOI:** 10.1101/127522

**Authors:** Anne Zirkel, Milos Nikolic, Konstantinos Sofiadis, Jan-Philipp Mallm, Lilija Brant, Christian Becker, Janine Altmüller, Julia Franzen, Mirjam Koker, Eduardo G Gusmao, Ivan G Costa, Roland T Ullrich, Wolfgang Wagner, Peter Nürnberg, Karsten Rippe, Argyris Papantonis

## Abstract

Ageing-relevant processes, like cellular senescence, are characterized by complex, often stochastic, events giving rise to heterogeneous cell populations. We hypothesized that entry into senescence of different primary human cells can be triggered by one early molecular event affecting the spatial organization of chromosomes. To test this, we combined whole-genome chromosome conformation capture, population and single-cell transcriptomics, super-resolution imaging, and functional analyses applied on proliferating and replicatively-senescent populations from three distinct human cell types. We found a number of genes involved in DNA conformation maintenance being suppressed upon senescence across cell types. Of these, the abundant high mobility group (HMG) B1 and B2 nuclear factors are quantitatively removed from cell nuclei before typical senescence markers appear, and mark a subset of topologically-associating domain (TAD) boundaries. Their loss coincides with obvious reorganization of chromatin interactions via the dramatic spatial clustering of CTCF foci. HMGB2 knock-down recapitulates this senescence-induced CTCF clustering, while also affecting insulation at TAD boundaries. We accordingly propose that HMGB-mediated deregulation of chromosome conformation constitutes a primer for the ensuing senescent program across cell types.

## Introduction

Since the original description of replicative senescence *in vitro* (Hayflick, 1965), *in vivo* implications of cellular senescence in development, wound healing, organismal ageing, and disease were revealed (Demaria et al., 2014; Munoz-Espin et al., 2013; Storer et al., 2013; van Deursen, 2014). In addition, the clearance of senescent cells in mouse models was recently shown to improve health-and lifespan (de Keizer, 2017). Senescence entry by primary mammalian cells is a result of integrated autocrine and paracrine signaling in the cell population (Acosta et al., 2013; Davalos et al., 2013; Hoare and Narita, 2016; Rodier et al., 2009) and leads to replicative arrest, and marked changes in gene expression, secretory activity, and chromatin structure (Ovadya and Krizhanovsky, 2014; Rai and Adams, 2013; Salama et al., 2014). As regards the chromatin landscape, changes like telomere shortening (Herbig et al., 2004), reorganization of heterochromatin and the lamina (Narita et al., 2006; Puvvula et al., 2014; Sadaie et al., 2013; Shah et al., 2013; Swanson et al., 2013; Zhang et al., 2005), activation of transposable elements (De Cecco et al., 2013), or epigenetic changes on histone tails and the primary DNA sequence occur (Cruickshanks et al., 2013; Franzen et al., 2017; Hanzelmann et al., 2015; Neyret-Kahn et al., 2013). These features of the transition into senescence render it a suitable model for studying the structure-to-function relationship of mammalian chromosomes – and chromosome conformation capture (3C) technologies now allow for this (Denker and de Laat, 2016).

Whole-genome conformation can be probed using Hi-C, a high throughput 3C variant that captures pairwise spatial interactions within and between chromosomes (Belton et al., 2012). As a result of Hi-C studies, we now understand that mammalian chromosomes are partitioned into large (multi-Mbp) active and inactive compartments (A-and B-compartments, respectively), which are in turn subdivided into consecutive topologically-associating domains (TADs; mostly sub-Mbp). TADs harbour a multitude of chromatin loops that tend to interact with one another more frequently than with loops in neighboring TADs (Dixon et al., 2012; Nora et al., 2012). This higher-order genomic organization is tightly linked to both the regulation of gene expression, but also to the transition through the cell cycle (Rowley and Corces, 2016).

To date, Hi-C has been applied to human fibroblasts inflicted by oncogene-induced senescence (OIS) or maintained for months in “deep” senescence. In OIS, a loss of local interactions within heterochromatin, including lamin-associated regions, was observed, and – in a presumed second step – the spatial clustering of some heterochromatic stretches that give rise to senescence-associated foci (Chandra et al., 2015). In “deep” senescence, short-range interactions were favored over longer-range ones resulting in the compaction of chromosomal arms, while centromeres tended to increase in volume (Criscione et al., 2016). What therefore remains an open question is whether spatial genomic reorganization is required for the very onset of senescence.

Here, we use three distinct human primary cell types from individual donors – umbilical vein endothelial cells (HUVEC), fetal lung fibroblasts (IMR90), and mesenchymal stromal cells (MSC) – in an effort to identify a shared regulatory backbone on the path towards replicative senescence. We combine high throughput genomics with super-resolution microscopy and single-cell sequencing to discover that non-histone proteins of the high mobility group B (HMGB) family are implicated in both the topological demarcation of TADs and the spatial rearrangement of CTCF-bound chromatin. HMGBs are abundant nuclear proteins characterized by the presence of a characteristic HMG-box DNA-binding domain; they are known for their ability to distort DNA via bending, looping or unwinding (Stros, 2010). This renders them important for gene transcription and nucleosome remodeling (Bonaldi et al., 2002; Das and Scovell, 2001; Laurent et al., 2010; Mitsouras et al., 2002; Monte et al., 2016; Redmond et al., 2015), as well as for genome integrity and recombination (Lee et al., 2010; Little et al., 2013; Polanska et al., 2012). However, HMGBs remain understudied as regards their global positioning on chromatin. Critically, HMGB1 and HMGB2 are depleted from cell nuclei in both senescent and ageing tissue (Abraham et al., 2013; Aird et al., 2016; Davalos et al., 2013; Taniguchi et al., 2009; Zhou et al., 2016), and are overexpressed across numerous types of cancer (http://www.cbioportal.org/; Gao et al., 2013). This work sheds new light onto the connection between the proliferative capacity of primary human cells, the maintenance of the genome’s spatial configuration and its transcriptional output.

## Results

### A shared regulatory backbone for replicative senescence across cell types

We hypothesized that there exists a common regulatory backbone for the entry into replicative senescence of different cell types. To assess this, we obtained cells from three disparate human cell types from individual donors (to gauge clonal variation): umbilical cord endothelial cells (HUVEC; mesodermal) from three donors, fetal lung fibroblasts (IMR90; endodermal) from two donors, and mesenchymal stromal cells (MSC; multipotent) from five donors. As the *in vitro* path towards senescence is characterized by heterogeneity (Smith and Whitney, 1980), we used both phenotypic and molecular markers to define the passage at which >60% (but not the bulk) of cells in these populations were entering senescence. Cells staining positive for β-galactosidase activity, showing significantly reduced proliferation in MTT assays, and are marked by changes in the methylation levels of six senescence-predictive CpGs (Franzen et al., 2017) were deemed as senescent (see Fig. 1A and **Supplemental Fig. S1A,B**). We should note here that, in the course of this *in vitro* passaging of cells, two donors (one from HUVECs and one from MSCs) displayed a limited number of population doublings until senescence and, despite their largely convergent gene expression profiles (*data not shown*), were excluded from all downstream analyses – this highlights the idiosyncratic nature of cellular ageing and its dependency on the donor’s genetic and physiological characteristics.

**Figure 1.**
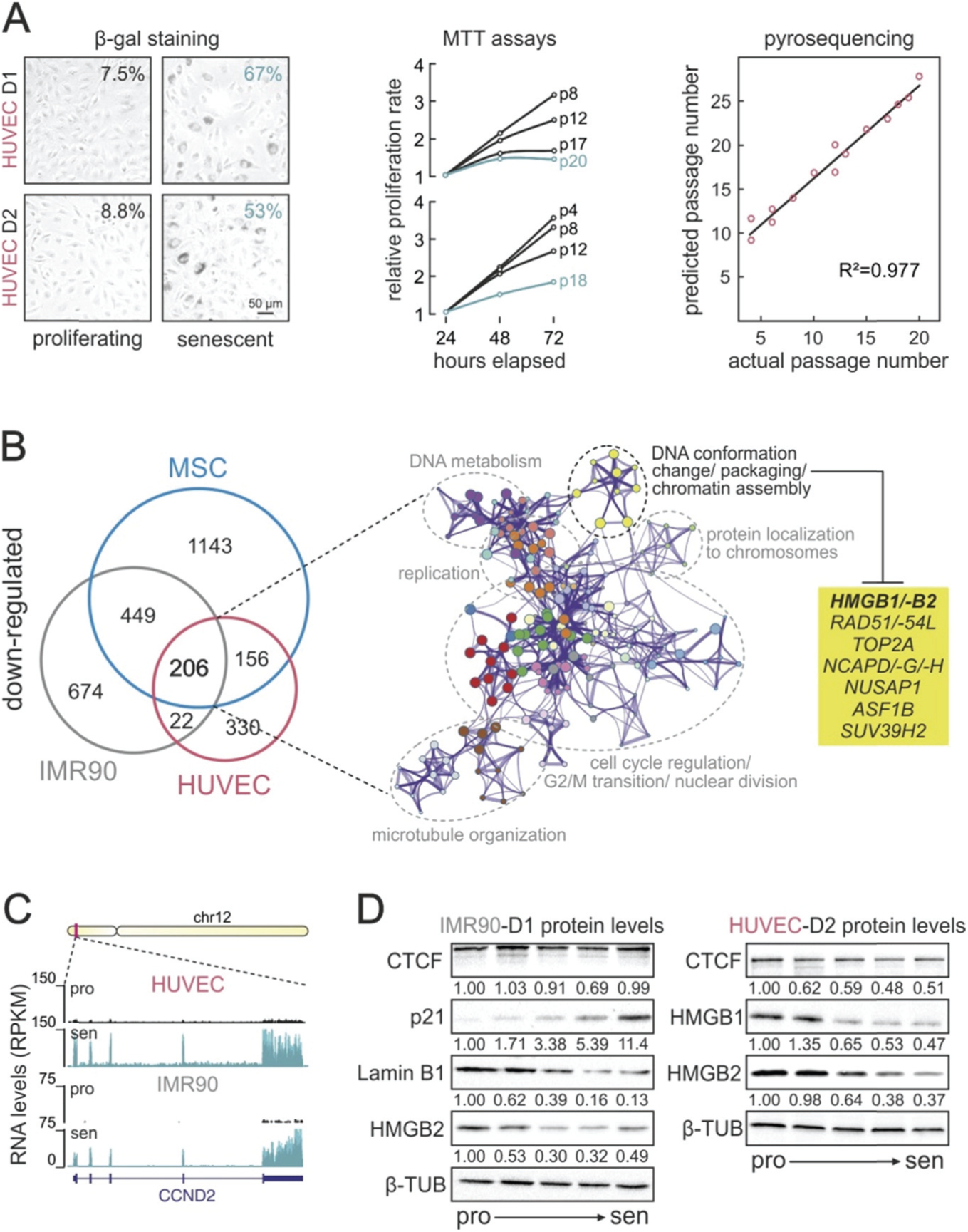
Senescence entry signals the suppression of specific pathways. (**A**) Senescence determination assays. *Left:* β-galactosidase staining of proliferating and senescent HUVECs from two individual donors (D1 and D2); percentages of β-gal positive cells are displayed. *Middle:* MTT proliferation assays over 72 hours for single-donor HUVECs at different passages; D1 cells in passage 20, and D2 cells in passage 18 were deemed as senescent (*green curves*). *Right:* Plot showing the correlation (R^2^) between actual and predicted passage number based on methylation levels of six CpGs measured by pyrosequencing. (**B**) Shared suppressed pathways in senescence. *Left:* Venn diagram showing genes down-regulated across all three cells types. *Right:* 206 shared down-regulated genes associate with the pathways in the displayed network. Nodes (*yellow*) containing genes involved in DNA conformation changes, DNA packaging, and chromatin assembly are highlighted, and selected ones are listed (*yellow box*). (**C**) Genome browser view of RNA-seq profiles along the *CCND2* locus on chromosome 12 that shows induced expression upon senescence in both HUVEC and IMR90 cells. (**D**) Protein levels of selected factors. Western blotting shows how the levels of HMGB1, HMGB2, CTCF, Lamin B1, and p21 change at five different passages in both HUVEC and IMR90 cells; numbers under each blot represent protein levels normalized to β-tubulin levels that serve as a control.

Early-passage (‘proliferating’) populations from all three cell types and donors were used to collect total RNA – and so were cell populations reaching replicative arrest (‘senescent’). Total RNA from all cell types/donors was depleted of rRNA species, enriched for poly-adenylated transcripts (with the exception of HUVEC samples) and sequenced to at least 50 million read pairs each. Following mapping to the reference genome (hg19), read counts were normalized using internal controls and/or spike-in cDNAs (Risso et al., 2014; **Supplemental Fig. S2**) to control for any differences in transcript abundance as a result of senescence. Then, differential gene expression analysis revealed different numbers of up-and down-regulated genes per cell type, which however grouped under similar gene ontology (GO) terms (**Supplemental Fig. S3A,B**). When the lists of up-regulated gene from all three cell types were intersected, 153 genes were shared and GO term analysis showed they were mostly involved the growth and p53 responses, and in extra-cellular matrix reorganization (**Supplemental Fig. S3C**). The 206 shared down-regulated genes associated with such processes as ‘cell cycle regulation’, ‘replication’, ‘DNA metabolism’, but critically also extended to include ‘DNA conformation change’, ‘chromatin organization/assembly’, and ‘DNA packaging’ (Fig. 1B). These last GO terms were due to the down-regulation in such genes as *HMGB1/B2, TOP2A, NCAPD/-G/-H, RAD51/-54L* or *ASF1*, and offered support to our hypothesis that spatial chromatin reorganization might occur early upon senescence entry. We validated these changes in gene expression by comparison to publicly-available data from proliferative/senescent IMR90 (Rai et al., 2014; Supplemental Fig. S3D), by RT-qPCR on selected targets in HUVEC (**Supplemental Fig. S3E**), as well as by isolating and analyzing nascent RNA from proliferative/senescent IMR90 via ‘factory RNA-seq’ (Melnik et al., 2016) to show that the core of the recorded changes is regulated at the level or transcription (Fig.1C and **Supplemental Fig. S4A-C**). Due to their *in vitro* ability to bend DNA, their depletion from ageing tissue *in vivo*, and their generally understudied role in chromatin looping, we focused on the HMGB1/B2 proteins and verified their suppression also at the protein levels by western blotting at different passages from IMR90 and HUVEC (Fig.1D).

### HMGB2 nuclear depletion upon senescence coincides with heterochromatic changes

Within the heterogeneous senescent populations we observed a characteristic increase in the size of cell nuclei, and these large (almost double in area) nuclei are typically in β-galactosidase positive cells (Fig.2A and **Supplemental Fig. S5A**). Focusing on larger-than-average nuclei, we were able to identify that senescent cells in particular show decreased levels of H3K27me3 (see also Shah et al., 2013), the main histone modification in facultative heterochromatin, and in the same nuclei increased levels of HP1α, a marker for constitutive heterochromatin (but no real formation of senescence-associated heterochromatin foci like those seen in OIS; (Narita et al., 2006; **Supplemental Fig. S5B**). This finding at the single-cell level is consistent with the senescent-induced drop of in the RNA levels of *EZH2*, the gene involved in H3K27me3 maintenance (**Supplemental Fig. S3E**). Although the increase in methylation of H3K9 was also seen at the population level using a multiplexed ELISA assay, the concomitant drop in H3K27me3 is only revealed via the analysis of single cells using microscopy (**Supplemental Fig. S5B-D**). In addition, histone acetylation levels decrease upon senescence entry as shown using H3K27ac and H4K16ac as markers (**Supplemental Fig. S5D,E**).

**Figure 2.**
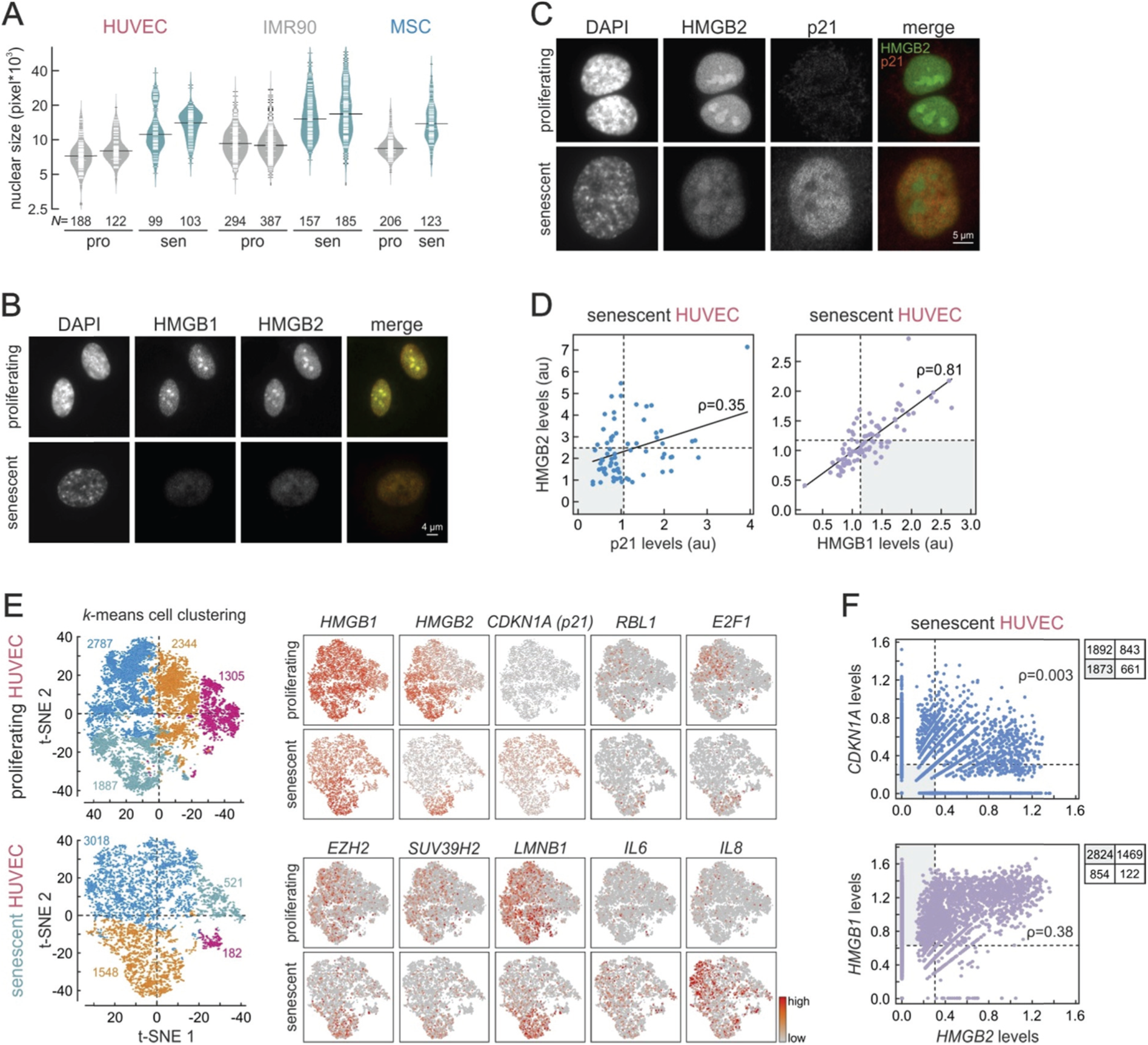
Entry into senescence is characterized by nuclear loss of HMGB1/B2. (**A**) Nuclear size increases upon senescence. Bean plots show the distribution of nuclear size in single-donor HUVEC, IMR90, and MSC population that were proliferating (‘pro’; *grey*) or senescent (‘sen’; *green*). The total number of nuclei measured (*N*) is indicated below. (**B**) Coordinated HMGB1/B2 loss in senescent cells. Widefield images of representative proliferating (*top*) and senescent HUVEC cells (*bottom*) immunostained for HMGB1 and HMGB2. (**C**) HMGB2 loss marks senescent cells. Widefield images of representative proliferating (*top*) and senescent HUVEC cells (*bottom*) immunostained for HMGB2 and p21. (**D**) Correlation between protein levels in single cells. Fluorescent levels of immunodetected HMGB2, HMGB1, and p21 from images like those in panels (B) and (C) were used to produce scatter plots. Median values minus one S.D. (*dotted lines*) and Pearson’s correlation coefficients (ρ) are shown. (**E**) Single-cell analysis of gene expression in HUVEC. *Left:* unbiased *k*-means clustering of proliferating (*top*) or senescent HUVEC (*bottom*); each cell is represented by a single dot, and the number of cells per cluster is indicated and colour-coded. *Right:* Plots showing the mRNA levels of ten senescence-regulated genes in the proliferating (*top rows*) and senescent cells (*bottom rows*). (**F**) Correlation between mRNA levels in single cells. Read counts for *HMGB2, HMGB1*, and *CDKN1A* from scRNA-seq were used to produce scatter plots. Median values minus one S.D. (*dotted lines*), Pearson’s correlation coefficients (ρ), and the number of cells in each quadrant are shown.

Next, we used super-resolution gSTED microscopy to investigate the subnuclear localization of HMGB1 and HMGB2 proteins; they are abundant nuclear components that mostly localize to low-DAPI signal regions (Supplemental Fig. S6A). Then, using widefield microscopy, HMGB1 and HMGB2 proteins were seen to be depleted from essentially every HUVEC, IMR90, and MSC senescent nucleus (also shown before; Aird et al., 2016; Davalos et al., 2013), and this loss is stronger in larger nuclei (Fig.2B and **Supplemental Fig. S6B**). Interestingly, whereas the events leading to the loss of HMGB1 and HMGB2 appear coordinated and correlated well in single cells (ρ=0.81), p21 upregulation – a hallmark of senescence establishment – correlates much less with HMGB2 nuclear loss (ρ=0.35; Fig. 2D). In fact, there exists a subpopulation of senescent HUVEC that have low HMGB2 levels, but have not yet gained in p21 levels, most probably indicative of HMGB1/B2 removal from nuclei being an early step on the path towards replicative senescence.

### Single-cell transcriptional profiling of senescent cells

In order to confirm that HMGB1/B2 are depleted from nuclei of cells entering senescence, before key replicative arrest markers are turned on, we performed single-cell mRNA sequencing on ˜8,300 proliferating (passage 4) and ˜5,200 senescent (passage 16) HUVEC using the DropSeq-based 10X Genomics platform. Following multiplexed sequencing, we generated at least 30,000 reads per each proliferating or senescent cell, and >2,500 unique gene transcripts were robustly captured per cell (see **Methods** for details). Then, we performed an unbiased clustering of cells per each condition based on single-cell expression profiles. This yielded four clusters for proliferating cells, and three for senescent ones (Fig.2E, *left*). Interestingly, the three major senescent-cell clusters largely reflect the relative distribution of *HMGB2* and *CDKN1A* (p21) expression. Much like at the protein level (see Fig.2D), low *HMGB2* levels tend to mark cells with medium-to-high *CDKN1A* levels (i.e., replicatively arrested cells) – but ˜1800 cells display low *HMGB2* without having significant *CDKN1A* expression. Similarly, >2,500 cells have strong *HMGB1* expression, but low *HMGB2* levels (Fig.2E,F). In addition, the *LMNB1* expression profile, as well as those of *EZH2* and *SUV39H2*, also appears to adequately describe the senescent subpopulations, indicative of the contribution of such epigenetic regulators in senescence entry. Conversely, cells in the senescent population lacking *HMGB2* expression are marked by high levels of the *IL6* and *IL8* SASP genes (Fig.2E). Thus, *HMGB2* suppression is documented as an early event that is decisive for HUVEC senescence entry.

### Senescence entry is marked by changes in 3D chromatin folding

RNA-seq and single-cell–derived data encouraged us to interrogate changes in whole-genome 3D chromatin folding that come upon senescence entry using Hi-C (Belton et al., 2012). We applied Hi-C under conditions that maintain nuclear integrity (Rao et al., 2014) using *HindIII* and sequencing to at least 400 million read pairs per each donor and condition. This allowed us to achieve a global, high-resolution understanding of spatial chromatin interactions in proliferating and senescent cells from all three cell types (down to a 20 kbp-resolution, much higher than what was previously achieved; (Criscione et al., 2016). Initially, we looked into how interchromosomal (*trans*) interactions change between proliferating and senescent cells; most chromosomes, in both HUVEC and IMR90 display decreased *trans*-interactions at the whole-chromosome level, although some chromosomes may behave in opposing ways between the two cell types (e.g., chromosomes 1 and 8; Fig. 3A). Then, we looked in detail into intrachromosomal (*cis*) interactions, and two key features mark the observed changes in interaction maps from proliferating and senescent cells. First, there is a striking increase in longer-range interactions upon senescence; second, we recorded stronger insulation in senescent cells, especially between larger, higher-order, domains (Fig. 3B and **Supplemental Fig. S7A**). This holds true for all three cell types and all chromosomes studied (see **Supplemental Fig. S8A-C**). When we subtract normalized interactions frequencies from 200 kbp-resolution maps, we found that subregions of the same chromosome are differentially-refolded upon senescence entry pending on the cell type investigated (Fig.3C and **Supplemental Fig. S7B**). However, interactions changes in individual donors are almost invariably of the same nature (Fig.3B and **Supplemental Fig. S7C**). Finally, in previously reported Hi-C data from fibroblast “deep” senescence, chromatin compaction was documented (Criscione et al., 2016). Our data agree with this observation, but with both longer-and shorter-range interactions e differentially emerging (Fig.3D, **Supplemental Figs S7D** and **S9**).

**Figure 3.**
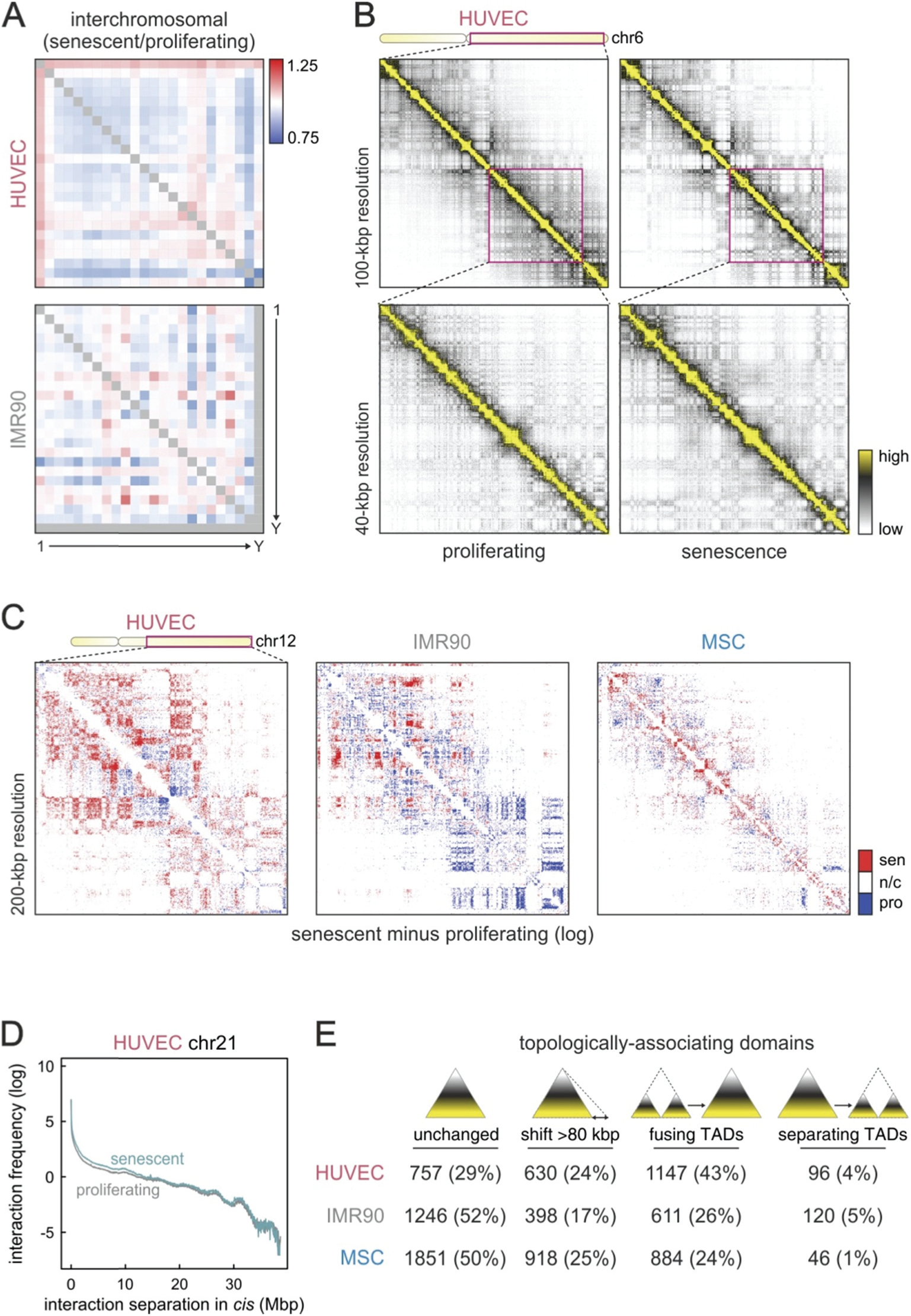
Senescence entry is accompanied by genome reorganization. (**A**) Changes in interchromosomal interactions. Heatmaps show increase/decrease (*red/blue*) in normalized Hi-C interaction frequencies between individual chromosomes from senescent versus proliferating HUVEC (*top*) and IMR90 (*bottom*). (**B**) Representative Hi-C heatmaps showing interaction frequencies along the long arm of chromosome 6 (*ideogram*) in proliferating and senescent HUVEC at 100- (*top row*) and 40-kbp resolution (*bottom row*). (**c**) Changes in intrachromosomal interactions. Heatmaps show binary increase/decrease (*red/blue*; see **Methods**) in normalized Hi-C interaction frequencies along the long arm of chromosome 12 (*ideogram*) from senescent versus proliferating HUVEC (*left*), IMR90 (*middle*), and MSC (*right*) at 200-kbp resolution. (**D**) Line plot showing normalized Hi-C interaction frequencies (log) for increasingly longer-range contacts along HUVEC chromosome 21 from proliferating (*grey*) and senescent cells (*green*). (**E**) Changes at the level of topologically-associating domains (TADs). The number (percentages in brackets) of TADs that remain unchanged, shift one boundary by >80 kbp, fuse into one larger TAD, or separate into smaller TADs upon senescence are listed for all three cell types studied here.

In a next step, we defined TADs in our Hi-C data from both conditions and all three cell types; to stay consistent with the original TAD definition (Dixon et al., 2012; Nora et al., 2012), we used an underlying 40-kbp resolution to do this and the TADtool suite (Kruse et al., 2016). Using the comparable calling parameters for both proliferating and senescent data from each cell type, this analysis returned ˜3,000 TADs in total for HUVEC and IMR90, and ˜3,500 for MSC in either state. By comparing TAD positions in proliferating and senescent cells of the same type, we found that ˜50% of them remain unchanged between the two states in IMR90 and MSC, ˜20% shift their boundaries by at least 80 kbp, and ˜25% fuse into larger TADs (Fig.3E). These changes were more drastic in HUVEC, with 30% TADs remaining unchanged, 24% shifting their boundaries, and 43% fusing into larger ones – which is also in line with the more widespread changes observed in HUVEC Hi-C maps (**Supplemental Fig. S7C**), and the greater TAD-boundary shifts recorded (**Supplemental Fig. S7E**).

### HMGB1 and HMGB2 mark gene promoters and topological domains boundaries

Our understanding of the roles of HMGB1/B2 *in vivo* remains incomplete, mostly due to the elusive nature of HMG-box factors in ChIP experiments (Redmond et al., 2015). Conventional crosslinking approaches not only fail to capture them bound on chromatin, but essentially repel them from it (Pallier et al., 2003; Teves et al., 2016). To overcome this, we applied a double crosslinking approach at a low temperature (see **Methods**), before immunoprecipitating HMGB1/B2-bound chromatin. Following ChIP-sequencing and data analysis, we identified 1574 robust HMGB2 peaks in proliferating HUVEC and 1188 in proliferating IMR90; we could also identify 391 peaks in a population of senescent IMR90 (for an example see Fig. 4A and **Supplemental Fig. S10A**). The majority of these peaks mark the promoters and bodies of active genes (Fig.4B) that are associated with processes relevant to senescence (e.g., ‘ECM organization’, ‘wound healing’ or ‘mitotic phase transition’; **Supplemental Fig. S10B**). It is noteworthy that at least twice as many genes bound by HMGB2 are up-rather than down-regulated in either cell type – and for those associated with the 131 peaks shared between HUVEC and IMR90, eight-fold more are upregulated and linked to such GO terms as ‘cell proliferation’, ‘regulation of proliferation’, and ‘cell ageing’ (Fig. 4C). On the other hand, there are 220 HMGB2 peaks that also remain bound in senescent IMR90. These peaks, just like the bulk of the 171 senescence-specific HMGB2 peaks, associate almost equally with up-and down-regulated genes, enriched for such GO terms as ‘M-phase regulation’, ‘mRNA surveillance’, and ‘post-transcriptional gene regulation’ (Fig.4D). However, we could not detect any HMGB2 ChIP signal enrichment upon senescence on SASP genes as previously reported (Supplemental Fig. S10C; Aird et al., 2016). Finally, we investigated DNase I-hypersensitive footprints under HMGB2 peaks for transcription factor (TF) motif enrichment. This analysis, after filtering for well-expressed TFs, returned a number of potentially co-bound factors, including structurally-important ones like CTCF or related to cell cycle regulation like members of the E2F family and FOXP1/P2 (**Supplemental Fig. S10D**). Then, looking into processes associated with up-/down-regulated such TFs, ‘negative RNA polymerase II regulation’, ‘inflammatory response’, ‘wound healing’ or ‘cell cycle regulation’ are enriched (**Supplemental Fig. S10D**, *bottom*).

**Figure 4.**
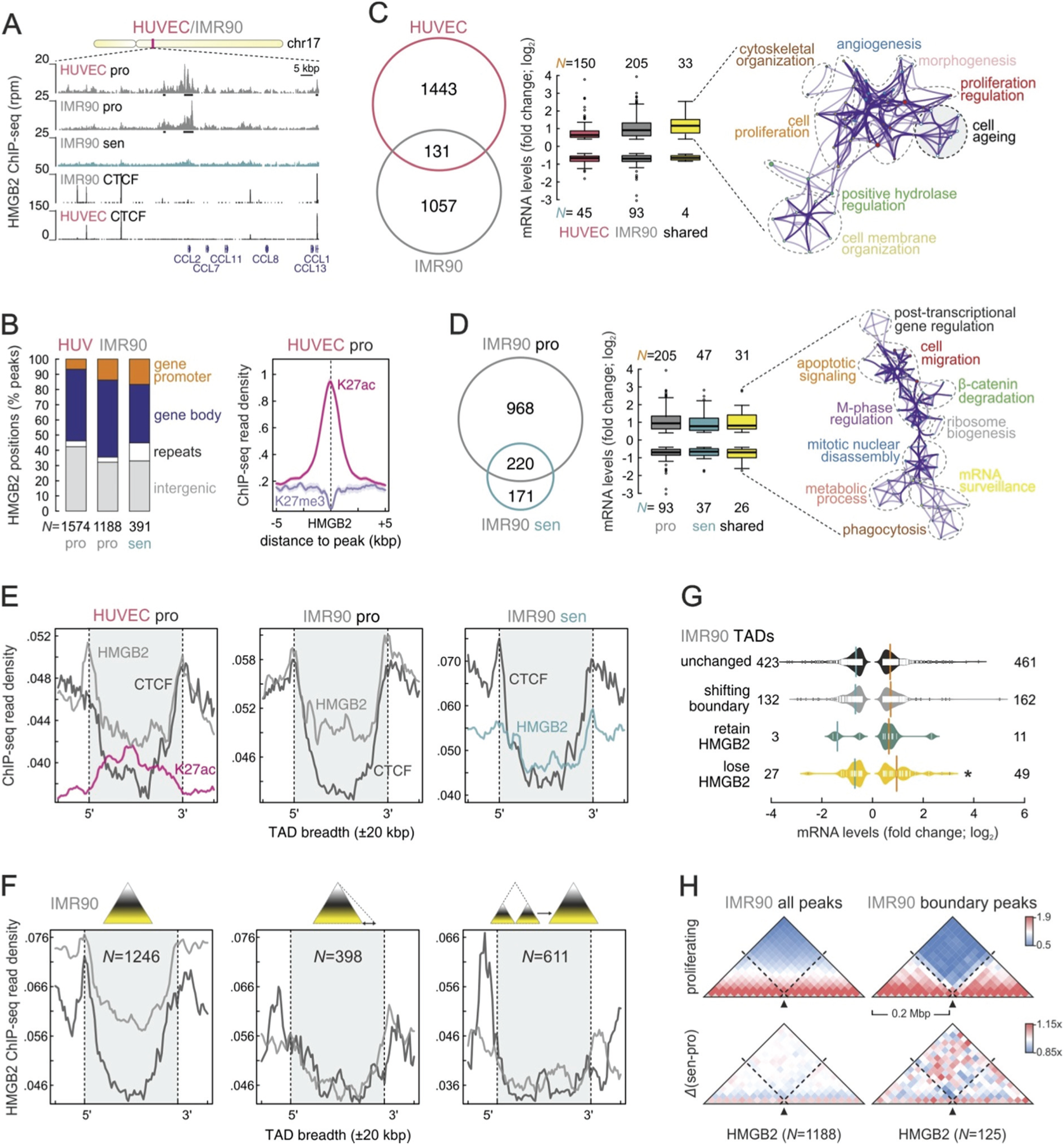
Chromatin-binding properties of HMGB2 change upon senescence entry. (**A**) Genome browser view of HMGB2 ChIP-seq profiles from proliferating (‘pro’) or senescent (‘sen’) HUVEC and IMR90 along a chromosome 12 locus harboring the *CCL* gene cluster; CTCF ChIP-seq profiles are also shown below. (**B**) Genomic distribution of HMGB2 peaks. *Left*: Bar plots showing the per cent of HMGB2 peaks from proliferating (‘pro’) or senescent (‘sen’) HUVEC and IMR90 overlapping gene promoters (*orange*), gene bodies (*blue*), repeats (*white*), or intergenic space (*grey*). The total number of peaks (*N*) per dataset is displayed below. *Right*: Line plot showing distribution of H3K27ac (*magenta*) and H3K27me3 ChIP-seq signal (*purple*) in the 10 kbp around all HUVEC HMGB2 peaks. (**C**) Shared HMGB2 peaks between proliferating HUVEC and IMR90. *Left*: Venn diagram showing peaks shared between the two cells types. *Middle*: Box plot showing the (log_2_) fold change in mRNA levels of differentially-regulated genes bound by HMGB2 in HUVEC, IMR90, or in both. The total number of up-/down-regulated genes (*N*) is displayed above/below. *Right*: 33 up-regulated genes bound by HMGB2 in both HUVEC and IMR90 associate with the GO terms/pathways in the displayed network. Nodes (*white*) containing genes involved in cellular ageing are highlighted. (**D**) Shared HMGB2 peaks between proliferating and senescent IMR90. *Left*: Venn diagram showing peaks shared between the two cell states. *Middle*: Box plot showing the (log_2_) fold change in mRNA levels of differentially-expressed genes bound by HMGB2 in proliferating (‘pro’), senescent (‘sen’) IMR90, or in both. The total number of up-/down-regulated genes (*N*) is displayed above/below. *Right*: 57 up-/down-regulated genes bound by HMGB2 in both cell states associate with the GO terms/pathways in the displayed network. (**E**) HMGB2 marks TAD boundaries. Meta-plots showing the distribution of HMGB2 ChIP-seq signal (*grey/green*) along all TADs (±20 kbp) called in proliferating HUVEC or proliferating/senescent IMR90. The distribution of CTCF (*black*) and H3K27ac (*magenta*) ChIP-seq signal serve as controls. (**F**) HMGB2 differentially marks changing and unchanging TAD boundaries. Meta-plots showing the distribution of HMGB2 (grey) and CTCF ChIP-seq signal (*black*) along IMR90 TADs (±20 kbp) that do not change (*left*), shift one boundary by >80 kbp (*middle*), or fuse into one larger TAD (*right*). The total number of TADs in each group (*N*) is also shown. (**G**) Differentially-expressed genes associated with TADs retaining/losing HMGB2. Bean plots showing the (log_2_) fold change in mRNA levels of differentially-expressed genes contained within (*from top to bottom*) TADs with unchanged or shifting boundaries (by > 80 kbp), and TADs that retain or lose HMGB2 from their boundaries in senescence. The total number of up-/down-regulated genes in each group (*N*) is also shown. *: significantly different mean; P<0.01, Wilcoxon-Mann-Whitney test. (**H**)Insulation by HMGB2 changes upon senescence. *Top*: Heatmaps showing normalized interaction frequencies in the 0.4 Mbp around all HMGB2 peaks (*left*) or around those at TAD boundaries (*right*) in proliferating IMR90. *Bottom*: Heatmaps showing interaction frequency changes in the 0.4 Mbp around all (*left*) or boundary HMGB2 peaks (*right*) between senescent/proliferating IMR90.

Using the same approach and analyses, we were able to investigate HMGB1 chromatin binding in proliferating HUVEC. We identified 810 robust HMGB1 peaks that do not overlap any HMGB2 peaks, and more than half of these are intergenic (**Supplemental Fig. S11A-C**). DNase I-footprints under HMGB1 peaks are enriched for factors like CTCF or E2F family members (**Supplemental Fig. S11D**). Similarly to HMGB2, HMGB1 binds twice as many up-rather than downregulated genes; the downregulated ones associate with such GO terms as ‘chromatin organization’, ‘cell cycle regulation’, ‘chromosome maintenance’, and ‘RNA polymerase II promoter regulation’ (**Supplemental Fig. S11E**).

Visual inspection of the HMGB1 and HMGB2 ChIP-seq data aligned to Hi-C data from the same cell types (**Supplemental Figs S11A** and **S12A**), revealed a potential correlation to positions of local insulation (often TAD boundaries). In a global analysis against TADs from proliferating HUVEC and IMR90, HMGB2 strongly marked their boundaries comparably to CTCF (Fig. 4E; this also applies to HMGB1 but to a lesser extent; **Supplemental Fig. S11F**). When we looked into HMGB2 ChIP-seq and TADs from senescent IMR90, boundary marking was weaker than that by CTCF; this held true when HMGB1/B2 ChIP-seq signal from proliferating HUVEC was plotted against TADs from senescent ones (Fig. 4E; **Supplemental Figs S12B** and **S11F**). On average, ˜10% of HMGB1/B2 peaks fall within 40 kbp of TAD boundaries in proliferating HUVEC or IMR90 (significantly more than what would be expected by chance; *P*<0.001, Fisher’s exact test), and we next asked how these boundaries behave in senescent cells. By plotting HMGB2 ChIP-seq signal against different TAD subgroups we find that HMGB2 marks the boundaries of both unchanged TADs, as well as of TADs with shifting boundaries (but not of fusing or separating TADs). Interestingly, those TADs that shift their boundaries have one boundary marked by HMGB2 and the other by CTCF (Fig. 4F and **Supplemental Fig. S12C**). Also, investigating the behaviour of genes included in different TAD subgroups, we found that those losing HMGB2 from their boundaries include more up-than downregulated genes, which also show stronger induction upon (Fig.4G). Lastly, in an effort to describe how HMGB2-bound TAD boundaries behave between senescent and proliferating cells, we plot Hi-C interactions in the 0.4 Mbp around HMGB2 boundary-peaks (compared to interactions arising around all peaks). These plots revealed that these boundaries essentially collapse upon senescence entry, which is not as prominent for all peaks (Fig.4H). A very similar effect was seen when this analysis was repeated for HMGB1 peaks (**Supplemental Fig. S11G**).

### HMGB2 depletion induces spatial CTCF clustering

The correlative analyses above are in support of a contribution by HMGB2 in the observed reorganization of 3D chromatin interactions upon entry into senescence. To establish this connection we performed *HMGB2* knock-down studies; as primary cells are difficult to transfect, we used self-delivering siRNA pools (see **Methods**) that efficiently deplete HMGB2 in both HUVEC and IMR90 (**Supplemental Fig. S13A**). Unlike the previously published effects of *HMGB1* knock-down (Davalos et al., 2013), within 72 h of treatment HMGB2 depletion only causes a mild increase in senescent cell numbers in HUVEC, does not severely impact DNA replication or the nuclear size of cells in the population (**Supplemental Fig. S13B,C**). Surprisingly, when we performed RNA-seq on two independent replicates of siRNA-treated HUVEC, we essentially did not detect genes strongly differentially-expressed (**Supplemental Fig. S13D**), which suggests that HMGB2 is not sufficient for robust transcriptional regulation. However, when we labeled nascent transcripts in living cells via a short (5-min) EU pulse and quantified resulting fluorescent EU-RNA levels on a widefield microscope, we saw a ˜30% drop in nascent RNA levels in HMGB2-knockdown cells compared to control ones. A similar decrease of ˜10% in nascent RNA production can also be seen in a senescent IMR90 population (Fig.5A).

**Figure 5.**
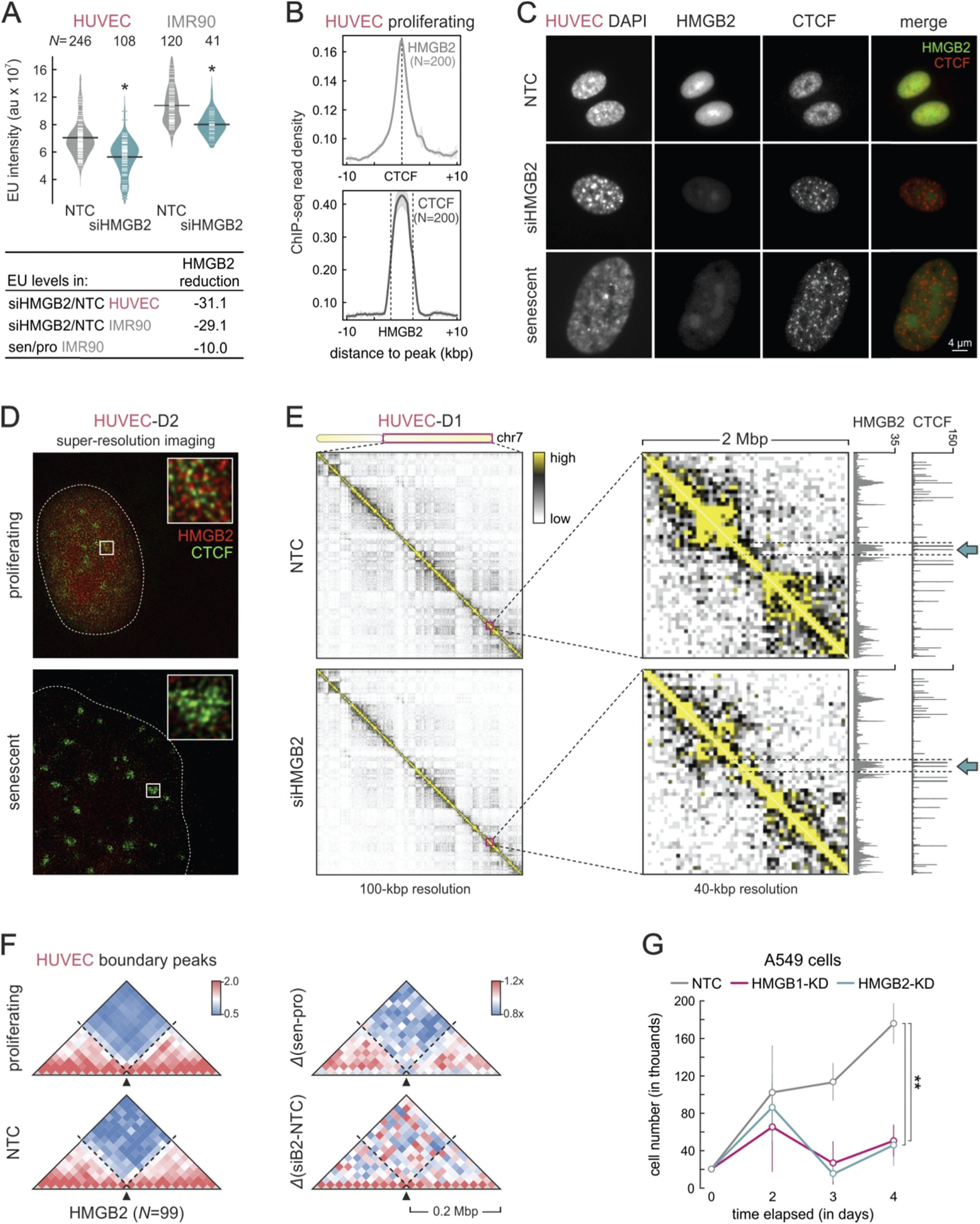
HMGB2 knock-down alters CTCF and chromatin 3D organization. (**A**) Nascent transcription decreases in the absence of HMGB2. *Top*: Bean plots showing a decrease in signal from EU-labeled nuclear RNA between control (‘NTC’) and knock-down (‘siHMGB2’) HUVEC or IMR90. *: significantly different; *P*<0.05, Wilcoxon-Mann-Whitney test. The total number of cells (*N*) measured in each condition is indicated above. *Bottom*: Summary of the measured reduction in nascent RNA levels from knock-down versus control HUVEC and IMR90, compared to that recorded in senescent versus proliferating IMR90 (normalized for the increase in nuclear size). (**B**) Relative positioning of HMGB2 and CTCF. Line plots showing HMGB2 ChIP-seq signal distribution in the 20 kbp around CTCF peaks (*top*) and vice versa (*bottom*), involving 200 peaks of each factor. (**C**) HMGB2 loss causes dramatic CTCF rearrangement. Widefield images of representative control (*top row*), knock-down (*middle row*), and senescent HUVEC (*bottom row*) immunostained for HMGB2 and CTCF. (**D**) Super-resolution view of senescence-induced CTCF clusters. gSTED images of representative proliferating (*top*) and senescent HUVEC (*bottom*) immunostained for HMGB2 and CTCF. (**E**) Representative Hi-C heatmaps showing interaction frequencies along the long arm of chromosome 7 (*ideogram*) in control (‘NTC’) and knock-down (‘siHMGB2’) HUVEC at 100- (*left*), and zoom-ins at 40-kbp resolution (*right*). HMGB2 and CTCF ChIP-seq data are aligned to the zoom-in heatmaps, and two typical regions with changed interactions are indicated (*arrows*). (**F**) HMGB2-marked TAD boundaries are remodeled. *Left*: Heatmaps showing normalized interaction frequencies in the 0.4 Mbp around HMGB2 peaks positioned at TAD boundaries in proliferating (*top*) or control HUVEC (‘NTC’; *bottom*). *Right*: Heatmaps showing changes in interaction frequencies in the 0.4 Mbp around HMGB2 boundary-peaks between senescent/proliferating (*top*) or siHMGB2/control HUVEC (*bottom*). (**G**)HMGB1/B2 knock-down attenuates proliferation of human lung adenocarcinoma (A549) cells. Line plot showing the number of living cells (mean ±S.D.; n=3) in culture at different times after transfection of siRNAs targeting HMGB1 (*magenta*) or HMGB2 (*green*); NTC transfections serve as a control (*grey*). **: significantly different mean; *P*<0.001, unpaired, two-tailed, Student’s t-test.

Based on the visual inspection of ChIP-seq data, proximity and overlap of HMGB2 and CTCF at domain boundaries was observed (**Supplemental Fig. S12A**). In HUVEC, there are 200 HMGB2 and CTCF peaks strongly overlapping one another – in IMR90, 172 CTCF peaks overlap HMGB2 ones in proliferating cells, and only 21 in senescent ones (Fig.5B and **Supplemental Fig. S13E**). Thus, we went on to image the CTCF distribution in control, *HMGB2*-knockdown, and proliferating/senescent HUVEC and IMR90. This revealed a dramatic spatial clustering of CTCF foci in senescent cells, which was faithfully recapitulated upon HMGB2 depletion (Fig.5C and **Supplemental S13F,G**). In fact, low HMGB2 nuclear levels predict the formation of these senescence-induced CTCF clusters (SICCs), and super-resolution imaging gives a detailed description of their multimeric structure. In proliferating HUVEC and IMR90 HMGB2 and CTCF foci are tightly interwoven, but in senescent/knockdown cells SICCs are strongly depleted of HMGB2 signal internally (Fig.5D and **Supplemental Fig. S14**). Importantly, SICCs do not overlap HP1α heterochromatin foci in senescence (**Supplemental Fig. S13H**). CTCF:CTCF loops from HUVEC and IMR90 (Rao et al., 2014) are not as strongly marked by HMGB2 as TAD boundaries are, and CTCF binding at standalone and HMGB2-cobound sites remains essentially unchanged between proliferating and senescent cells (**Supplemental Fig. S13I,J**).

We next treated a larger cell number with the siRNA pool to knock-down *HMGB2* and performed Hi-C. Interaction heatmaps, already at a 100-kbp resolution, show obvious differences between control and knockdown cells – and inspection of 40-kbp resolution maps in respect to HMGB2 binding peaks revealed strong contact remodeling at these sites in particular (Fig.5E and **Supplemental Fig. S15A**). Plotting Hi-C interactions in the 0.4 Mbp around HMGB2 boundary-peaks from knockdown versus control cells, we could observe a stronger collapse of insulation (Fig.5F), which also held true when looking into interactions around all HMGB2 peaks (**Supplemental Fig. S15B**). Strikingly, interactions around CTCF sites from HUVECs also change to reflect SICC formation, and the effect is stronger once all CTCF peaks are considered – but, surprisingly not when HMGB2 and CTCF co-bound peaks are interrogated (**Supplemental Fig. S15C**). Finally, when looking into changes at the TAD level upon knock-down, these are largely like those in senescent HUVEC (**Supplemental Fig. S15D,E**).

### HMGB1 and B2 control the proliferative capacity of lung adenocarcinoma cells

HMGB1 and HMGB2 are overexpressed across a large number of cancer types (TCGA data; http://www.cbioportal.org/; Gao et al., 2013), and HMGB1 overexpression has been directly linked to the proliferative capacity and metastasis of malignant cells (Tang et al., 2010) and of lung adenocarcinoma in particular (Feng et al., 2016). Hence, we sought to investigate how HMGB2-mediated deregulation of chromosome organization might affect lung adenocarcinoma patient lines. Under our dual crosslinking conditions, HMGB2 appears associated with chromatin throughout cell division (**Supplemental Fig. S16A**; also shown by overexpression and live-cell imaging by Pallier et al., 2003). We speculated that it might be topologically-bookmarking chromatin in anticipation of the exit from mitosis, and thus knocking-down HMGB2 in cancer cells where it is highly expressed (**Supplemental Fig. S16B**) should affect proliferation rates. Indeed, when three different lung adenocarcinoma lines – A549, NCI-H1650, and NCI-H1975 – were transfected with the same siRNA pool used for HUVEC/IMR90, severe growth arrest was observed within ˜48 h (similar to that achieved by *HMGB1* knock-down in the same lines; Fig. 5G and **Supplemental Fig. S16C**). However, RT-qPCR analysis in these knockdown cells revealed that *HMGB2* suppression does not affect *HMGB1* levels, but *HMGB1* knockdown also turns off *HMGB2* expression. This points to a hierarchical relationship between the two factors, which in turn differentially affect such effector genes as *HDAC9, RBL1, CTCF*, or *TP53* (while *LMNB1, EZH2*, and *SUV39H2* are equally affected in both knockdowns; **Supplemental Fig. S16D**).

## Discussion

Proliferating cells have mechanisms in place that preserve their inherent chromatin states and, presumably, their higher-order architecture in conjunction with DNA replication and cell division. For example, 3D folding of chromosomes appears to gradually dissolve towards mitosis, before being reinstated after cell division concludes (Naumova et al., 2013). However, it is not clear how this might be achieved in proliferatively-arrested cells. Conversely, tumour cells exhibit dramatically enhanced proliferative capacities and should therefore have hyper-activated chromatin preservation mechanisms in conjunction with cell division.

We hypothesized that, despite their different developmental origins and functional characters, the entry into senescence by different cell types is governed by a shared regulatory backbone – presumably involving an early step of chromatin reorganization. Using endothelial cells, lung fibroblasts, and mesenchymal stromal multipotent cells, we were able to assign this presumed commonality to genes associated with DNA conformation and its maintenance. Of this subset of genes, HMGB1 and HMGB2 stood out because of their inherent ability to distort chromatin and their high nuclear abundance (Stros, 2010). Both HMGB1 and HMGB2 are strongly depleted from cell nuclei upon senescence entry, and for HMGB1 it has been demonstrated that this can induce cell cycle arrest in fibroblasts (Davalos et al., 2013). Although HMGB2 knock-down only mildly affected DNA replication, gene expression, or nuclear size, it does cause a widespread reduction of nascent RNA levels – a feature of senescent cells.

Critically, though, a subset of HMGB2 (and HMGB1) binding-peaks mark the boundaries between TADs/sub-TADs, where spatial chromatin interactions are insulated. At these sites, boundaries most often coincide with an active gene promoter, which is consistent with an early observation that active protein-coding and tRNA genes are found at TAD boundaries (Dixon et al., 2012). We recently proposed that transcription acts as a force partitioning mammalian genomes (Zirkel and Papantonis, 2014); looking into changes at HMGB-marked boundaries, it is evident that the change in activity of the underlying gene (mostly up-regulation, if only mild) coincides with spatial contact reshuffling upon senescence. Moreover, in *HMGB2*-knockdown HUVEC, these same boundaries display an even more evident collapse. Notably, both senescent and HMGB2-knockdown cells are characterized by the dramatic clustering of CTCF foci into SICCs – this, in conjunction with the heterochromatic remodeling seen in senescence cells (but not upon *HMGB2* knock-down), give rise to the strong reorganization seen in Hi-C data. Notably, such a stress-induced clustering was very recently observed *in vivo* in mice with deficient DNA repair mechanism (Chatzinikolaou et al., 2017). Inevitably, the differential regulation of additional chromatin-binding factors, like topoisomerase II (Uuskula-Reimand et al., 2016) or HMGB1, will also contribute to chromosome refolding in senescence. Hence, replicative senescence constitutes a highly suitable model system for perturbing and studying the structure-to-function relationship of human chromosomes.

We also hypothesized that the somewhat stochastic propagation of senescence in the cell population, via both autocrine and paracrine signaling, is triggered by an early molecular event. The single-cell RNA-seq and microscopy analyses presented here are in support of this. We can clearly detect a subset of cells in the senescent populations that have lost HMGB2 from their nuclei, but still have some HMGB1 (of which a fraction will be on its way to secretion; (Bonaldi et al., 2003)) and no detectable p21 expression. Similarly, *IL6* and *IL8* expression, which is central to the SASP activation, is mostly confined to HMGB2-depleted cells, whereas for *LMNB1* the converse applies.

Then, we propose that there exists a hierarchy of events that essentially begin by the deregulation of higher-order chromatin organization via HMGB2 nuclear eviction and constitute a primer for the ensuing senescent program in different cell types. It is noteworthy that HMGB1 and HMGB2 do not display redundant functions; they bind non-overlapping genomic regions, appear to associate with discrete co-bound TF subsets, and exhibit different temporal behaviour upon senescence entry. Then, also given their different spatiotemporal expression in development (Kang et al., 2014; Ronfani et al., 2001), HMGB1 and HMGB2 should perhaps also be decoupled as regards to their *in vivo* effects in respect to senescence (see also Bagherpoor et al., 2017). HMGB2 acts as a rheostat of topological insulation and thus affect global transcription competency. Finally, the fact that we can detect HMGB2 bound to mitotic chromatin (in a manner similar to another HMG-box factor, Sox2; Teves et al., 2016) implies a possible role for it in mitotic bookmarking of these same positions, thus connecting proliferative capacity to topological demarcation. Although further work is needed to dissect this possibility, it may explain the inability of HMGB2-overexpressing lung adenocarcinoma cells to propagate in the persistent absence the factor.

## Methods

### Cell culture and senescence markers

HUVECs from single, apparently healthy, donors (passage 2; Lonza Inc.) were continuously passaged to replicative exhaustion in complete Endopan-2 supplemented with 2% FBS under 5% CO_2_. Cells were constantly seeded at ˜10,000 cells/cm^2^, except late passages that were seeded at ˜20,000 cells/cm^2^. Single-donor IMR90s (I90-10 and −79, passage 5; Coriell Biorepository) were continuously passaged to replicative exhaustion in MEM (M4655, Sigma-Aldrich) supplemented with non-essential amino acids and 20% FBS under 5% CO_2_. Mesenchymal stromal cells (MSCs) were isolated at the Aachen Medical School, and cultured to senescence as previously described (Franzen et al., 2017).

Cell proliferation was monitored by MTT assays at different passages. In brief, ˜5,000 cells are seeded in 96-well format plates in quadruplicates. On the next day, the medium is replaced with 100 μl fresh medium plus 10 μl of a 12 μM MTT stock solution (Invitrogen), and cells are incubated at 37°C for 4 h. Subsequently, all but 25 μl of the medium is removed from the wells, and formazan dissolved in 50 μl DMSO, mixed thoroughly and incubated at 37°C for 10 min. Samples are then mixed again and absorbance read at 530 nm. Measurements are taken at 24, 48 and 72 h post-seeding, background subtracted, and normalized to the 24 h time point. Senescence-associated β-galactosidase assay (Cell Signaling) was performed according to the manufacturer’s instructions to evaluate the fraction of senescent cells at different passages. Finally, DNA methylation at six selected CpG islands (Franzen et al., 2017) was measured by isolating genomic DNA at the different cell states and performing targeted pyrosequencing (Cygenia GmbH).

### Immunofluorescence and image analysis

Cells were grown on coverslips from the stage indicated; they were fixed in 4% PFA/PBS for 15 min at room temperature. After washing once in PBS, cells were permeabilized with 0.5% Triton-X/PBS for 5 min at room temperature. Blocking with 1% BSA/PBS for 1h was followed by incubation with the indicated primary antibodies for 1-2 h. Cells were washed twice with PBS for 5 min before incubating with secondary antibodies for 1 h at room temperature. Nuclei were stained with DAPI (Sigma-Aldrich) for 5 min, washed, and coverslips mounted onto slides in Prolong Gold Antifade (Invitrogen). For image acquisition, a widefield Leica DMI 6000B with a HCX PL APO 63x/1.40 (Oil) objective was used; confocal and super-resolution images were acquired on a Leica TCS SP8 gSTED microscope with a 100x/1.40 (Oil) STED Orange objective. Deconvolution of the super-resolution images was performed using the Huygens software from Scientific Volume Imaging.

For image analysis, the NuclearParticleDetector2D of the MiToBo software (from version 1.4.3; available at (http://www2.informatik.uni-halle.de/agprbio/mitobo/index.php/Main_Page) was used. Measurements of nuclear immunofluorescence signal were automatically generated using a mask drawn on DAPI staining to define nuclear bounds. Background subtractions were then used to precisely determine the mean intensity per area of each immunodetected protein. The following primary antibodies were used: α-CTCF (1:1000, 61311, Active Motif), α-p21 (1:500, ab184640, Abcam), α-HMGB2 (1:1000, ab67282, Abcam), α-HMGB2 (1:1000, 12248-3D2, Sigma), α-HMGB1 (1:1000, ab190377, Abcam), α-Tubulin (1:1000, T0198, Sigma Aldrich), HP1α (1:1000, 39977, Active Motif), H3K9me3 (1:200, 39286, Active Motif), H3K27me3 (1:1000, C15410069, Diagenode), H3K27ac (1:500, C15410174, Diagenode), H4K16ac (1:500, 61529, Active Motif). For STED microscopy 2C Pack STED 775 IR-R- (1:2000, 2-0032-052-6, Abberior) secondary antibodies were used.

### Genome-wide chromosome conformation capture (Hi-C) and analysis

For Hi-C studies, 25-35 million proliferating/senescent cells were crosslinked by adding 1% PFA (Electron Microscopy Sciences) for 10 min at room temperature with subsequent quenching (125 mM glycine), and gently scraped off the plates on ice. Then, Hi-C was performed as previously described (Belton et al., 2012) under conditions where nuclei remained largely intact. In brief, cells were lysed (10 mM Tris-HCl pH 8.0, 10 mM NaCl, 0.4% NP40, 1x protease inhibitor cocktail) twice for 20 min on ice, washed with 1x NEBuffer 2 twice, and distributed into 4-5 tubes. Following SDS treatment (0.1%) for 10 min at 65°C and quenching with Triton-X for 15 min at 37°C, chromatin was digested with 600 units of *Hind*III overnight at 37°C under constant agitation. Next day, digestion was extended for another hour with an additional 600 units of *Hind*III for 1-2 hours. DNA overhangs were filled-in using a biotin-dCTP containing mix (1.5μl dATP, 1.5 μl dGTP, 1.5 μl dTTP, 37.7 μl biotin-dCTP, each 10 mM stock; Invitrogen) in the presence of 1.5 μl 50 U/μl Klenow (NEB) for 45min at 37°C, and all enzymes are inactivated by adding 86 μl 10% SDS to all tubes and incubating for 30min at 65°C. Next, each individual mixture was transferred into a 15 ml tube, incubated for 10-20min on ice to quench SDS, and a mixture containing 745 μl of 10% Triton-X, 745 μl 10x ligation buffer (500 mM Tris-HCl pH 7.5, 100 mM MgCl_2_, 100 mM DTT), 80 μl of a 10 mg/ml BSA stock, 80 μl of 100 mM ATP and 5.96 ml milliQ water is added to each tube, and blunt-end ligation initiated by adding 15 μl of 5 U/μl T4 DNA ligase (Invitrogen) and incubated for 6 hours at 16°C. Crosslinks were then reversed and proteins degraded by adding 35 μl of 15mg/ml proteinase K and incubated overnight at 65°C. Next day, an additional 35 μl of proteinase K was added for 2 hours at 65°C, mixtures were cooled to room temperature, transferred to 50 ml conical tubes, and DNA was extracted twice with phenol:chloroform (ph 8.0) and ethanol precipitated. The resulting pellets were dissolved in 450 μl 1x TE, transferred to 1.7ml tubes, DNA was re-extracted twice, precipitated, pellets were washed once in 70% ethanol, air-dried, and resuspended in 25 μl 1x TE. Subsequently, three tubes, each with ˜5 μg of the Hi-C template, were incubated for 2 h at 12°C in the presence of 1 μl 10 mg/ml BSA, 10 μl 10x NEBuffer 2, 1 μl 10 mM dATP, 1 μl 10 mM dGTP (for *Nco*I samples dCTP was used instead) and 5 units of T4 DNA polymerase (NEB) in a total volume of 100 μl to remove biotin-16-dCTP at non-ligated DNA ends. Reactions were stopped by adding 2 μl of 0.5 M EDTA pH 8.0, and the DNA was subsequently purified by phenol:chloroform extraction followed by ethanol precipitation. DNA pellets were resuspended and pooled in a total of 100μl of milliQ water, and residual salts were removed via Amicon 30K columns using three washing steps (1x 200 μl and 2x 100 μl milliQ water). Finally, DNA was sheared to a size of 300-600 bp on a Bioruptor Plus (Diagenode; 2x 10 cycles of 30 sec *on* and 30 sec *off*, at the highest power setting).

For IMR90 cells, the procedure was slightly modified to follow the *in situ* Hi-C protocol (Rao et al. 2014). Approx. 20 million cells were lysed twice in 900 μl lysis buffer (10 mM Tris-HCl pH 8.0, 10 mM NaCl, 0.4% NP-40, 1x protease inhibitor cocktail) for 15 min on ice, nuclei were pelleted and washed once in 1x NEBuffer 2, distributed into 4 tubes, resuspended in 200 μl 0.5% SDS/NEBuffer 2, and incubated for 10 min at 62°C. After repelleting, SDS-treated nuclei were resuspended in 150μl 1% Triton-X/NEBuffer 2, and digested overnight using 400 units of *Hind*III at 37°C with constant agitation. Next day, another 400 units of *Hind*III were added for another 1-2 h at 37°C. Following heat inactivation for 20 min at 62°C, nuclei were pelleted by centrifugation and resuspend in 256 μl NEBuffer 2. DNA ends were then biotinylated by adding 50 μl of a biotin-dCTP-Klenow mix (37.5μl of 0.4 mM biotin-dCTP, 1.5μl of 10 mM dTTP, dATP, and dGTP, and 1.5μl of 50 U/μl Klenow) to each tube and incubating for 1 h at 37°C. Subsequently, 900 μl of ligation mix (603 μl H_2_O, 240 μl 5x Invitrogen ligation buffer, 30 μl of 10% Triton-X, 12 μl of 10 mg/ml BSA, and 15 μl ligase) was added to each sample, mixed, and incubated for 6 h at 16°C; tubes were inverted frequently. Nuclei were next pelleted and a volume of 900 μl was removed. Following reversal of crosslinking and proteinase K addition, DNA is phenol:chloroform extracted and ethanol precipitated; all subsequent steps were performed as described above.

Hi-C DNA was used as template for adding sequencing linkers via only 10 PCR cycles. Then, Hi-C libraries were sequenced to generate at least 400 million paired-end reads (50-75 bp in length) per each donor/condition in a HiSeq4000 platform (Illumina). Raw reads were mapped to the human reference genome (hg19) iteratively (to ensure maximum recovery of uniquely mapped pairs) using BWA (Li and Durbin, 2010), duplicates were removed (http://picard.sourceforge.net/), and the output converted into BEDPE format (Quinlan and Hall, 2010). Next, custom R scripts were used to bin the genome into non-overlapping bins (typically 10-kbp ones), assign reads to bins, remove read pairs not representing valid interactions, and to normalize read counts to library size. Finally, the HiTC Bioconductor package (Servant et al., 2012) was used to annotate and correct the matrices for biases in genomic features (Yaffe and Tanay, 2011) and to visualize 2D heat maps at different resolutions. For subtracted Hi-C maps, we determined bins under a particular cutoff (typically 5 rpm) and set them to 0, while all othesr were set to 1; then, matrices were subtracted to give a binary appreciation of the interaction changes between the different states. For plotting insulation heatmaps, normalized interactions values in the twenty 20-kbp bins around each HMGB1/B2 peak were added up, normalized to the median value in each bin and plotted. All R scripts used here are available on request.

### RNA isolation, sequencing, and analysis

Cell of different types/conditions were harvested in Trizol LS (Life Technologies) and total RNA was isolated and DNase I-treated using the Direct-zol RNA miniprep kit (Zymo Research). Following, rRNA-depletion using the RiboZero Gold kit (Illumina), barcoded cDNA libraries were generated using the TruSeq RNA library kit (Illumina) with (IMR90, MSC) or without selection on poly(dT) beads (HUVEC); in addition some IMR90 and MSC libraries were spiked using a synthetic ERCC Spike-In mix (Thermo Fischer Scientific) to facilitate normalization. The resulting libraries were paired-end sequenced to at least 50 million read pairs on a HiSeq4000 platform (Illumina), and raw reads were mapped to the human reference genome (hg19) and analyzed via the QuickNGS pipeline (Wagle et al., 2015) to obtain read counts. Then, read counts from different single-donors/conditions were normalized using the RUVSeq package (Risso et al. 2014). Please note that, in our hands, normalization using either synthetic spike-in controls or the intrinsic ‘top quantile’ function of the RUVseq package is essentially interchangeable. Finally, for analysis of nascent RNA in IMR90 the ‘factory RNA-seq’ approach was applied on ˜5 million proliferating or senescent cells (Melnik et al., 2016), RNA was isolated and sequenced as above, and intronic read counts were obtained and differentially analyzed for the two conditions using the iRNAseq package (Madsen et al., 2015). All box/bean plots were plotted using the BoxPlotR online (http://shiny.chemgrid.org/boxplotr/), and GO term/pathway enrichment analyses using the Metascape interface (http://metascape.org/gp/index.html; (Tripathi et al., 2015)). Differentially-regulated genes per each cell type are listed in **Supplemental Table S1**.

### Chromatin immunoprecipitation (ChIP), sequencing, and analysis

For each batch of ChIP experiments approx. 25 million cells, cultured to >80% confluence in 15-cm dishes, were crosslinked in 15 mM EGS/PBS (ethylene glycol bis(succinimidyl succinate); Thermo) for 20 min at room temperature, followed by fixation for 40 min at 4°C in 1% PFA. From this point onwards, cells were processed using the ChIP-IT High Sensitivity kit (Active motif) as per manufacturer’s instructions. Chromatin was sheared to 200-500 bp fragments on a Bioruptor Plus (Diagenode; 2x 9 cycles of 30 sec *on* and 30 sec *off*, at the highest power setting), and immuno-precipitation was carried out by adding 4 μg of the appropriate antiserum (HMGB1: PCRP-HMGB1-4F10-s, Developmental Studies Hybridoma Bank; HMGB2: ab 67282, Abcam; CTCF: 61311, Active motif) to approx. 30 μg of chromatin and incubating on a rotator overnight at 4°C in the presence of protease inhibitors. Following addition of protein A/G agarose beads and washing, DNA was purified using the ChIP DNA Clean & Concentrator kit (Zymo Research) and used in qPCR or next-generation sequencing on a HiSeq4000 platform (Illumina). Where ChIP-seq was performed, at least 35 million reads were obtained and the respective ‘input’ sample was also sequenced. Raw sequencing reads (typically 100 bp-long) were mapped to the reference human genome (hg19) using BWA (Li and Durbin, 2010), and the resulting .BAM files were processed via the MACS2 software (Zhang et al., 2008) to identify signal enrichment over input. Thresholded HMGB1/B2 ChIP-seq peaks per each cell type are listed in **Supplemental Table S2**, and oligonucleotides used as primers in ChIP-qPCR listed in **Supplemental Tables S3** and **S4**. The .BAM files were also used in ngs.plot (Shen et al., 2014) for plotting ChIP-seq coverage over particular genomic positions for different conditions/cell types. Finally, transcription factor recognition motif enrichments within DHS footprints under HMGB1/B2 ChIP-seq peaks were calculated using the Regulatory Genomics Toolbox (Gusmao et al., 2014).

### Western blotting

For assessing protein abundance at the different cell states, approx. 4×10^6^ cells were gently scraped off 15-cm dishes, and pelleted for 5 min at 600 x *g*. The supernatant was discarded, and the pellet resuspended in 100 μl of ice-cold RIPA lysis buffer (20 mM Tris-HCl pH 7.5, 150 mM NaCl, 1 mM EDTA pH 8.0, 1 mM EGTA pH 8.0, 1% NP-40, 1% sodium deoxycholate) containing 1x protease inhibitor cocktail (Roche), incubated for 20 min on ice, and centrifuged for 15 min at >20,000 x *g* to pellet cell debris and collect the supernatant. The concentration of the nuclear extracts was determined using the Pierce BCA Protein Assay Kit (Thermo Fisher Scientific), before extracts were aliquoted and stored at −70°C to be used for western blotting. Resolved proteins were detected using the following antisera (and dilutions): α-CTCF (1:2000, 61311, Active Motif) α-Lamin B1 (1:2000, ab16048, Abcam), α-p21 (1:1000, ab184640, Abcam), α-HMGB2 (1:2000, ab67282, Abcam), α-HMGB1 (1:1000, ab190377, Abcam), α-Rad21 (1:1000, ab992, Abcam), α-Tubulin (1:2000, T0198, Sigma Aldrich), and visualized using the Pierce SuperSignal West Pico ECL kit (Thermo Fisher Scientific).

### EdU and EU labeling of nucleic acids

Nascent DNA synthesis was monitored by EdU incorporation and subsequent labelling with the Click-iT^^®^^ chemistry (Click-iT EdU Imaging Kit; Invitrogen). In brief, cells were incubated in 10 μM EdU for 7 h, fixed using 3.7% PFA/PBS for 15 min at room temperature, permeabilized, and labeled as per manufacturer’s instructions. Similarly, nascent RNA synthesis was monitored by incorporating EU in transcripts by incubating cells in 1 mM of 5-ethynyl uridine for exactly 5 min at 37°C. Cells were then fixed, permeabilized, and labeled via Click-iT chemistry as above. Before imaging on a widefield Leica microscope as described above, cells were also immunostained for HMGB2, stained with DAPI, and mounted onto glass slides.

### Single-cell mRNA sequencing and analysis

Freshly-frozen early- (p. 4) and late-passage HUVEC (p. 16) were thawed, washed once in warm PBS, and subjected immediately to encapsulation in oil droplets. Each droplet contained beads carrying barcoded oligos for cDNA synthesis that allow both cell barcoding and unique transcript identification on a 10X Chromium Genomics Drop-seq platform as per manufacturer’s instructions. This way 8,323 early-and 5,269 late-passage cells were processed (with a 0.8% chance of capturing a cell duplet). Samples from each condition were pooled and sequenced separately in two HiSeq2500 lanes each (Illumina) yielding ˜30,000 reads per early-and ˜47,000 reads per late-passage cell. This accounts for a >50% sequencing saturation, and returned >2,500 and >3,000 robustly captured transcripts for early- and late-passage cells, respectively. Raw sequencing data were processed and visualized using default settings in the Cell Ranger suite provided by the manufacturer.

### siRNA-mediated HMGB2 knock-down

HUVEC or IMR90 were seeded at 20,000 cells/cm^2^ one day before transfection. An Accell-siRNA pool (Dharmacon) against *HMGB2*, plus a non-targeting control (NTC; fluorescencently-tagged to allow transfection efficiency to be monitored for each experiment), were added to the cells at a final concentration of 1 μM. Knock-down efficiency and its downstream effects were evaluated at 72 h after transfection using RT-qPCR and/or immunofluorescence assays. For HUVEC Hi-C, cells were seeded the same way but in 2x15cm plates per condition. The cells were crosslinked at 72h post transfection and processed as described for IMR90 and MSC Hi-C. Due to the lower cell number (˜5 million cells) permeabilized nuclei were kept in a single tube for downstream processes.

### Statistical analyses

P-values associated with Student’s t-tests were calculated using the GraphPad online software (http://graphpad.com/), those associated with the Wilcoxon-Mann-Whitney test using the EDISON-WMW tool (Marx et al., 2016), while Pearson’s correlation coefficients (ρ) using the function embedded in the MS Excel suite (Microsoft).

### Data availability

All NGS data generated here are available at the NCBI GEO repository (*accession number pending*).

## Supplemental information

This paper is accompanied by **Supplemental Figs S1-S16** and **Supplemental Tables S1-S4**.

## Acknowledgements

We would like to thank George Garinis and members of the Papantonis lab for critical reading of the manuscript, and the CECAD microscopy facility for help with gSTED imaging. This work was supported by Interdisciplinary Center for Clinical Research (IZKF) funding (to IGC), by the Else Kröner Fresenius Stiftung (to WW and AP), by a DFG basic module grant (RI 1283 14/1; to KR and AP), and by CMMC core funding, a UoC Advanced Researcher Grant and the Fritz Thyssen Stiftung (all awarded to AP).

## Author contributions

AZ, MN, and AP conceived research; AZ, KS, and LB performed experiments; JPM and KR sequenced and analyzed single-cell RNA-seq libraries; CB, JA, and PN assisted with Hi-C library preparation and performed all high throughput sequencing; MN analyzed RNA-seq, ChIP-seq, and Hi-C data; JF and WW sequenced and analyzed DNA methylation data; EGG and IGC performed DHS-centered motif analysis of ChIP-seq data; AZ, MN, and AP wrote the manuscript with input from all co-authors.

